# TRIM25 inhibits Influenza A infection by destabilising its mRNA and is redundant for the RIG-I pathway

**DOI:** 10.1101/2021.09.13.460052

**Authors:** Nila Roy Choudhury, Gregory Heikel, Ivan Trus, Rute Maria Dos Santos Pinto, Maryia Trubitsyna, Eleanor Gaunt, Paul Digard, Gracjan Michlewski

**Author notes:** MRC-University of Glasgow Centre for Virus Research, Glasgow, United Kingdom. Correspondence should be addressed to G.M.

## Abstract

The E3 ubiquitin ligase TRIM25 is a key factor in the innate immune response to RNA viruses. TRIM25 has been shown to play a role in the <u>r</u>etinoic-acid-<u>i</u>nducible <u>g</u>ene-<u>1</u> (RIG-I) pathway, which triggers expression of type 1 interferons upon viral infection. We and others have recently shown that TRIM25 is an RNA-binding protein, however not much is known about the RNA-binding roles of TRIM25 in the innate immune response to RNA viruses. Here, we demonstrate that influenza <u>A</u> <u>v</u>irus (IAV A/PR/8/34_NS1(R38K41A)) infection is inhibited by TRIM25. Surprisingly, host RNA-binding deficient mutant TRIM25ΔRBD and TRIM25ΔRING, which lack E3 ubiquitin ligase activity rescued IAV inhibition in TRIM25 knock-out cells. Furthermore, we show that in human cultured cells activation of the RIG-I/interferon type 1 pathway mediated by an IAV-derived 5’-triphosphate RNA does not require TRIM25 activity. Additionally, knocking out TRIM25 does not affect the activity of the IAV polymerase. We present new evidence that TRIM25 restricts IAV by directly binding to and destabilising its mRNAs. Finally, we show that direct tethering of TRIM25 to RNA is sufficient to downregulate the targeted RNA. In summary, our results uncover a novel mechanism that TRIM25 uses to inhibit IAV infection and regulate RNA metabolism.

## Introduction

Influenza A virus (IAV) is a ubiquitous human pathogen that causes seasonal epidemics and sporadic pandemics, the most recent of which was in 2009. Seasonal IAV kills up to 500,000 people annually, thereby causing a significant global socioeconomic burden^1^. Vaccines are updated yearly to target contemporary strains, but this relies on making predictions about an unpredictable virus. Alternative interventions targeting host cell factors are attracting increasing interest to overcome the issues forthcoming due to high viral diversity. To identify such targets, it is essential to understand how viruses interact with host cell factors.

The innate immune system is the body’s first line of defence against viral infection. Viruses may be detected by the host cell, triggering signalling cascades that result in the expression of interferon proteins and antiviral interferon-stimulated genes ^2,3^. E3 ubiquitin ligase TRIM25 (a member of the tripartite motif (TRIM) family of proteins) has emerged as a key factor in triggering the innate immune response to RNA viruses ^4,5^. All TRIMs have in common their amino-terminal tripartite domain arrangement — RING–Bbox1/2–coiled coil (CC) — yet differ in their C-terminal domains, informing their characterisation into several subtypes. TRIM25 has been shown to play a role in the retinoic-acid-inducible gene-1 (RIG-I) pathway, which triggers expression of type I interferons upon viral infection ^6^. RIG-I binds to RNA molecules with a 5’-triphosphate (5’ppp) moiety that are produced during viral replication ^7–9^. This provokes a RIG-I conformational change allowing TRIM25 multimers to bind to and ubiquitinate the RIG-I tandem caspase recruitment domains (2CARD), thus triggering a series of events that culminate in the phosphorylation of IRF-3, IRF-7 and NF-κB. These transcription factors then translocate to the nucleus and induce type I interferon expression, leading to an innate immune response ^10–12^. There is also evidence that RIG-I signalling can be activated by unanchored K63-linked polyubiquitin chains that are generated by TRIM25 ^13^. Recent data however suggests that TRIM25 may not have a key role in RIG-I activation as previously proposed ^14,15^.

We and others have revealed that TRIM25 is an RNA-binding protein ^16–20^. TRIM25 binds host RNAs through a novel <u>R</u>NA-<u>b</u>inding <u>d</u>omain (RBD) residing in its C-terminal PRY/SPRY region ^21^. RNA-binding appears to be crucial for its E3 ubiquitin ligase activity as deletion of the RNA-binding domain reduces TRIM25 ubiquitination in vitro ^21^. Specific sequence or conformational RNA motifs which attract TRIM25 binding have not been identified.

Consistent with a role in antiviral sensing, TRIM25 is known to be inhibited by several RNA viruses to dampen the innate immune response ^19,22–24^. For IAV, this is via the non-structural protein 1 (NS1). NS1 directly binds to the coiled-coil domain of TRIM25, disrupting its dimerization and consequently, its ubiquitin ligase activity ^22,25^. However, although NS1 proteins from several IAV strains bind to TRIM25, not all of these are capable of inhibiting phosphorylation of IRF-3 (presumably following RIG-I activation) upon virus infection ^26–28^, indicating that NS1 may be targeting an alternative function of TRIM25. Additionally, NS1 can also bind to and antagonise TRIM25 function in chicken cells, where no RIG-I orthologue has been identified, hinting at a RIG-I-independent antiviral function for TRIM25 ^26^. Indeed, recent work has indicated that TRIM25 can restrict IAV by binding to viral ribonucleoproteins (RNPs) and inhibiting viral mRNA chain elongation ^29^. This interaction is RNA-dependent, involving direct binding of TRIM25 to the IAV genomic vRNA, but detailed map and functions of TRIM25-RNA binding to IAV are unknown.

Here, we confirm that in human HEK293 cells, IAV replication is sensitive to TRIM25 in the presence of a functional NS1 protein mutant. However, inhibition of viral replication by TRIM25 did not require TRIM25 E3 ubiquitin ligase activity or the PRY/SPRY RNA binding domain nor did it correlate with the ability of virally derived 5’-pppRNA to stimulate IRF-3 phosphorylation or IFN expression. Using CLIP-seq we show that TRIM25 binds preferentially to positive strand IAV RNAs. Furthermore, the inhibitory function of TRIM25 did not appear to work by direct inhibition of viral transcription. This suggests that TRIM25 is not necessary for RIG-I activation in human cultured cells and uses alternative, RIG-I-independent mechanisms in restricting IAV infection. Indeed, we identify a novel mechanism whereby TRIM25 affects the stability of IAV mRNAs. Crucially, direct tethering of TRIM25-MS2 fusion protein to MS2 stem loops integrated into the 3’UTR of the luciferase mRNA significantly reduced RNA levels. These results shed light on TRIM25 in the innate immune response to RNA viruses and present a new molecular mechanism by which it controls the stability of the RNA.

## Results

### TRIM25 inhibits IAV replication irrespective of its RBD or RING domains

We have previously described the generation of a HeLa TRIM25 knock-out (HeLaΔTRIM25) cell line through CRISPR/Cas9-mediated gene editing ^21^. The same technique was used to generate a HEK293 TRIM25 knock-out cell line (TRIM25 KO) ^30^. This cell line contains a flippase recognition target (FRT) site to allow stable integration of a gene of choice into the genome using the Flp-In recombinase ^31^. Codon optimised TRIM25 WT, TRIM25ΔRBD (lacking RNA-binding residues 470–508 located in the PRY/SPRY domain) and TRIM25ΔRING, which does not have E3 ligase activity, were integrated into the genome at this site, resulting in stable expression at levels close to endogenous TRIM25 in HEK293 wild-type (WT) cells (Fig. 1A). These cell lines were infected with the IAV A/PR/8/34 (PR8) strain, either WT virus or a virus with R38A/K41A (R38K41A) mutations in NS1 that render it unable to inhibit TRIM25 and is defective in blocking the RIG-I pathway ^22,32–35^. Cells were infected at a low multiplicity of infection and viral replication was assessed with the endpoint dilution assay (Fig. 1B). In WT HEK293 cells WT PR8 virus replicated better than the PR8 R38K41A mutant, presumably due to the presence of a fully active NS1 protein inhibiting TRIM25. In contrast, in the HEK293 TRIM25 KO cells the PR8 WT and R38K41A mutant grew to similar titers. Surprisingly, however, the latter phenotype was not only rescued by integrated WT TRIM25 but also by TRIM25ΔRBD and partially by the integration of the ubiquitination mutant TRIM25ΔRING (Fig. 1B). TRIM25 KO cells were growing noticeably slower than WT cells, thus cell-to-cell comparison with regards to IAV replication was not possible. Together, these results suggested that in TRIM25 anti-viral activity against IAV is not entirely dependent on its E3 ligase activity nor due to its fragment of the PRY/SPRY domain which binds RNA.

**Figure 1.**
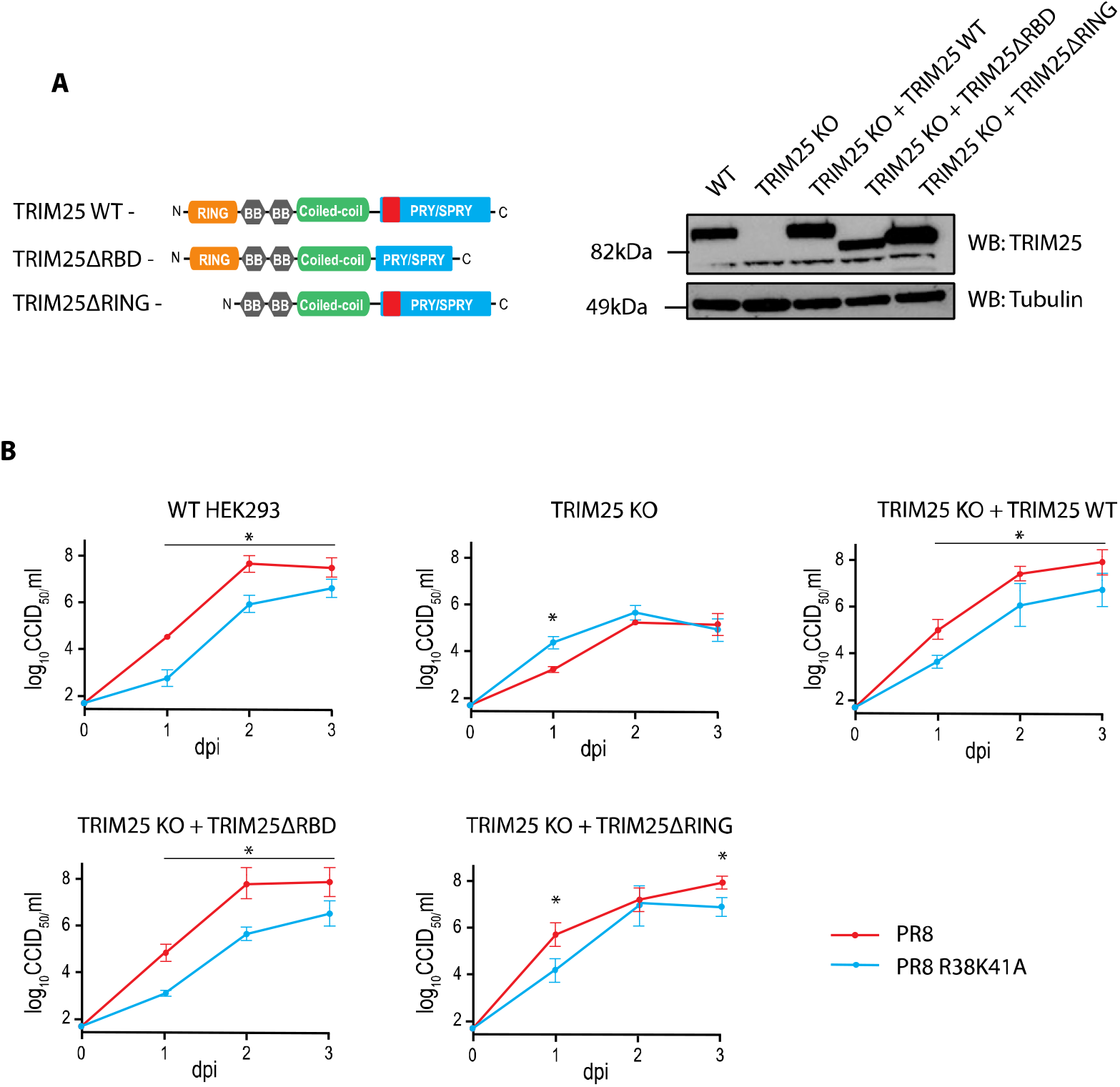
IAV replication defect in TRIM25 KO HEK293 cells can be rescued with RNA binding or ubiquitination deficient TRIM25 mutants. (A) TRIM25 WT, TRIM25ΔRBD or TRIM25ΔRING were re-integrated into HEK293ΔTRIM25 cells using the Flp-In recombinase system and levels of the proteins were compared to WT cells by western blot. Domain architecture of wild-type TRIM25 and deletion mutants. Relative position of host RNA-binding domain (RBD) is shown in red. (B) Cell lines were infected with IAV PR8 WT or PR8 NS1 R38K41A (MOI=0.0001) and virus titers were assessed by the endpoint dilution assay. Whiskers represent the standard error of the mean (SE) from three technical replicates. Dpi—days post-inoculation. The asterisk (*) indicates p < 0.05 in two-way ANOVA with Sidak’s post-test.

### Both TRIM25 and TRIM25ΔRBD bind to positive IAV RNAs during infection

To test if TRIM25 binds IAV RNAs and to determine if the RNA-binding deficient TRIM25ΔRBD was still able to bind viral RNAs, we performed a CLIP-seq as published previously^21^. HEK293ΔTRIM25 cells with integrated T7-tagged TRIM25 WT or TRIM25ΔRBD (Fig. 2A) were infected with the IAV PR8 R38K41A strain at an MOI of 5 for 1 or 6 hours followed by UV crosslinking (Fig. 2B). While after 1h little interaction was observed, following 6 hours of infection both TRIM25 WT and TRIM25ΔRBD was shown to interact with positive-strand IAV RNAs (Fig. 2B). This could be either mRNAs or the replication intermediate cRNAs. Both TRIM25 WT and TRIM25ΔRBD displayed interaction with positive RNAs derived from IAV segments. The interaction with negative strand RNAs was at the threshold of signal detection by the CLIP-seq. Segments 4, 5, 7 and 8 presented more pronounced TRIM25 peaks than segments 1-3 and 6. This is most likely because segments 4, 5, 7 and 8 are expressed at much higher levels upon IAV infection than segments 1-3 and 6 ^36^. Importantly, TRIM25ΔRBD does not bind to the majority of host RNAs as compared to WT TRIM25, as seen on SDS PAGE with 5’-end labelled total RNA crosslinked and immunoprecipitated with anti-T7 Ab (Fig. 2C).

**Figure 2.**
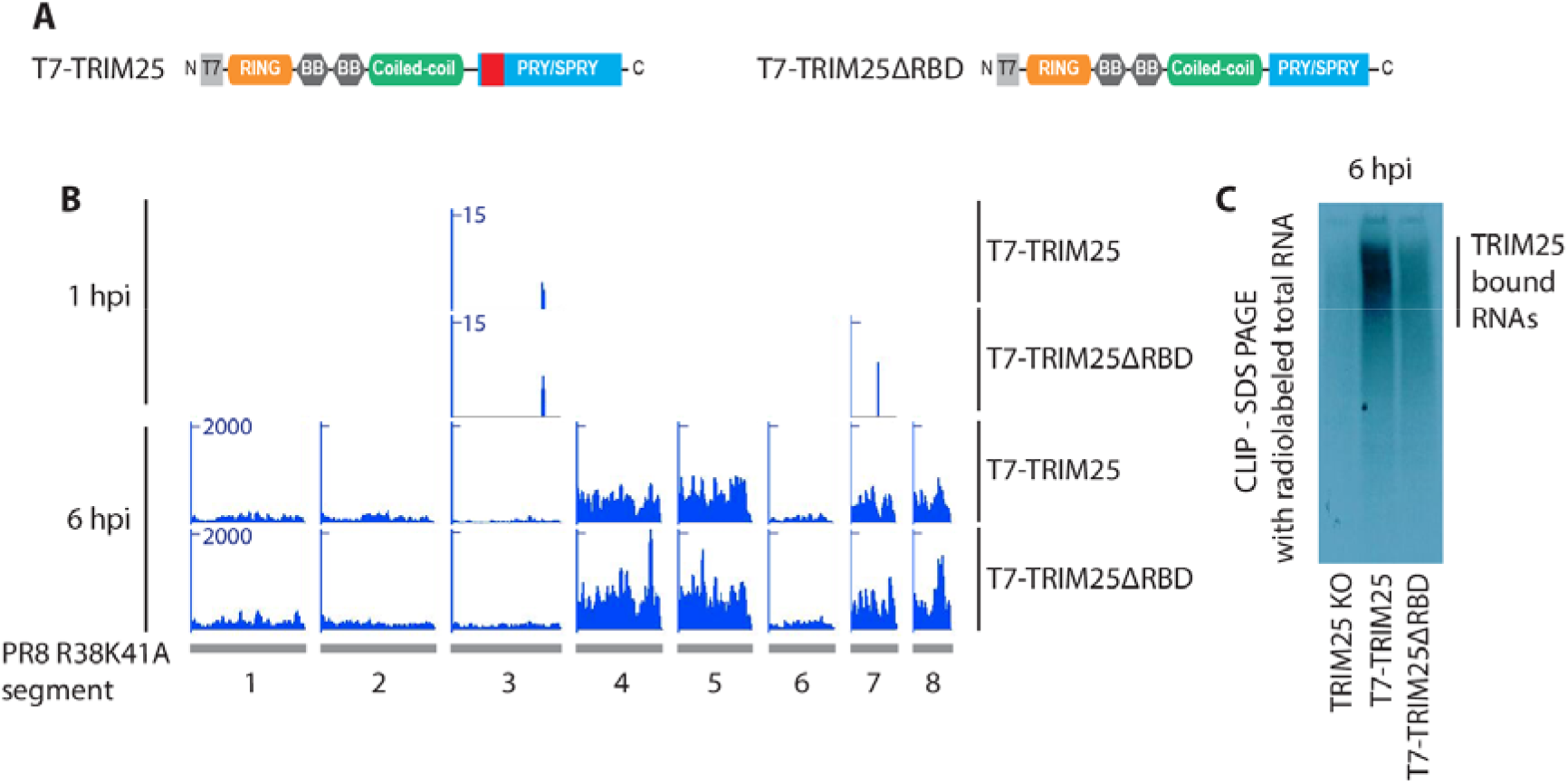
TRIM25 binds positive viral RNA. (A) Domain architecture of wild-type TRIM25 and TRIM25ΔRBD tagged with T7. (B) CLIP-seq analysis of TRIM25 binding to IAV RNA shows that both WT TRIM25 and TRIM25ΔRBD can bind to positive-sense Influenza RNA. TRIM25 binds moderately to positive-sense IAV RNA (1 hour post infection – 1 hpi) and abundantly to viral mRNAs or cRNA (6 hours post infection – 6 hpi). (C) TRIM25ΔRBD does not bind to majority of host RNAs as seen on the SDS PAGE with 5’-end labeled, total RNA crosslinked and immunoprecipitated with the anti-T7 Ab.

Segment 7 had the highest number of Transcripts Per Million (TPM) at 6 hours post infection both in original (TPM=145177) and TRIM25ΔRBD HEK cells (TPM=79554). Surprisingly, allocation of these reads measured upon infection moderately correlated (Spearman’s ρ=0.43) with the GpC (guanine-phosphate-cytosine) dinucleotide frequency in the IAV genome in the original HEK cells (Supplementary Figure 1, Supplementary Table 1), but not in modified TRIM25ΔRBD HEK cells (ρ=0.11). On the contrary a lack of correlation was identified for the mirrored CpG dinucleotide both in original (ρ=−0.09) and modified HEK cells (ρ=0.06). Other IAV segments had lower number of mapped reads and subsequently a slightly lower correlation level was identified in the original HEK cells at 6 hours post infection for GpC dinucleotide (ρ=0.33 for segment 5 and ρ=0.41 for segment 8). This is in line with our previous data suggesting that TRIM25 prefers binding to RNA motifs rich in G and C ^21^.

These results for the first time demonstrate a full spectrum of TRIM25 binding to IAV-derived RNAs and show that it preferentially binds positive-strand IAV RNAs. This could be because of the lower abundance of negative-strand RNAs, and the fact that the negative-strand RNA is interacting with IAV’s NP as well as other viral and host factors. Finally, binding of TRIM25ΔRBD uncovers additional modes of RNA recognition and explains why the mutant restricts IAV infection to the same levels as WT TRIM25.

### TRIM25 is not required for the IFN response to 5’ppp-RNA

To assess the ability of the HEK293 cell lines to signal through RIG-I, they were transfected with a synthetic 5’ppp panhandle RNA (3p-hpRNA) derived from segment 8 of IAV which is known to induce signalling through RIG-I ^37^. Cells were transfected with 3p-hpRNA and incubated for 6 hours before levels of IRF-3 phosphorylation were analysed by western blot (Fig. 3A). Production of IFNα/β under the same conditions was also assayed using the HEK-Blue bioassay system (Fig. 3B). Neither assay showed a difference in activation of RIG-I signalling between the cell lines with wild type TRIM25, without TRIM25 or with stably integrated TRIM25ΔRING or TRIM25ΔRBD. In addition, the activity of the IFNβ promoter in response to 3p-hpRNA transfection was assayed using a Firefly luciferase reporter construct. 3p-hpRNA was co-transfected into cells alongside the IFNβ promoter-Firefly plasmid and a plasmid constitutively expressing Renilla luciferase as a loading/transfection control and 24 hours later levels of luciferase were assayed (Fig. 3C). This assay also showed no significant difference in IFNβ promoter activity in cells that did not express TRIM25 compared to those that expressed wild type or mutant TRIM25. Taken together, these results strongly suggest that in HEK293 cells TRIM25 is not necessary for activation of RIG-I signalling and that the loss of virus restriction in HEK293 TRIM25 KO cells cannot be explained by differences in IRF-3 activation.

**Figure 3.**
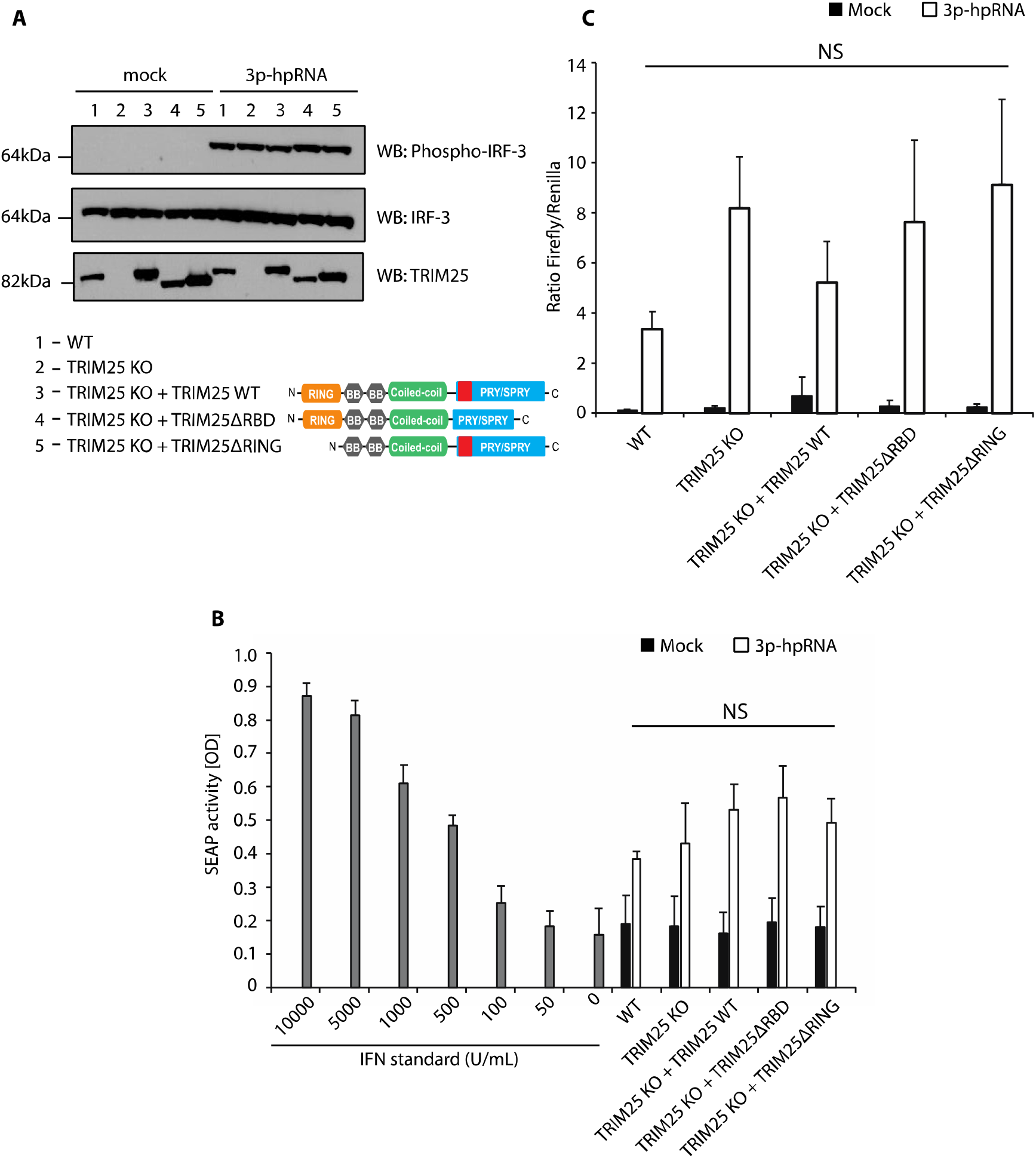
Deletion of TRIM25 from HEK293 cells does not affect activation of RIG-I signalling. (A) HEK293 cell lines were transfected with 100 ng/mL of the IAV-derived synthetic RNA 3p-hpRNA and incubated for 6 hours. Levels of IRF-3 phosphorylation were assayed by western blot. Domain architecture of wild-type TRIM25 and deletion mutants. Relative position of host RNA-binding domain (RBD) is shown in red. (B) HEK-Blue assay measuring secreted IFNα/β from each cell line 6 hours after transfection with 100 ng/mL of 3p-hpRNA. The means and SDs of three independent experiments are shown. Statistical significance was calculated using one-way ANOVA with Tukey’s post-test. (C) Dualluciferase assay measuring induction of the IFNβ promoter in each cell line 24 hours after transfection with 100 ng/mL of 3p-hpRNA. Renilla luciferase was used as a loading and transfection control. The means and SDs of three independent experiments are shown. Statistical significance was calculated using one-way ANOVA with Tukey’s post-test.

### RIG-I activation is reduced in the absence of TRIM25 in MEF, but not HeLa, cell lines

To test whether this result was limited to HEK293 cells, RIG-I activation by 3p-hpRNA was assayed in other TRIM25 knock-out cell lines. Previously generated HeLa TRIM25 KO cells ^21^, in addition to HeLa wild type cells, were transfected with 3p-hpRNA and levels of phospho-IRF-3 were assayed by western blot after 6 hours (Fig. 4A). As seen in HEK293 cells, there was no difference in activation of RIG-I/IRF-3 signalling between WT and HeLa TRIM25 KO cells. If anything, IRF-3 phosphorylation was more pronounced in HeLa TRIM25 KO cells. When levels of secreted IFNα/β were analysed from the same experiment using the HEK Blue assay, we did not observe statistically significant stimulation of IFNα/β (Fig. 4B). Finally, WT and TRIM25 KO mouse embryonic fibroblasts (MEFs - a gift from Prof. Jae Jung, University of Southern California ^38^) were analysed for levels of phospho-IRF-3 in response to transfection of 3p-hpRNA after 6 hours (Fig. 4C). MEF WT cells showed higher levels of phospho-IRF-3 than MEF TRIM25 KO cells, indicating that in these cells TRIM25 is important for activation of RIG-I signalling. Of note, our total IRF-3 antibody did not react with mouse IRF-3 and therefore DHX9 was used as a loading control.

**Figure 4.**
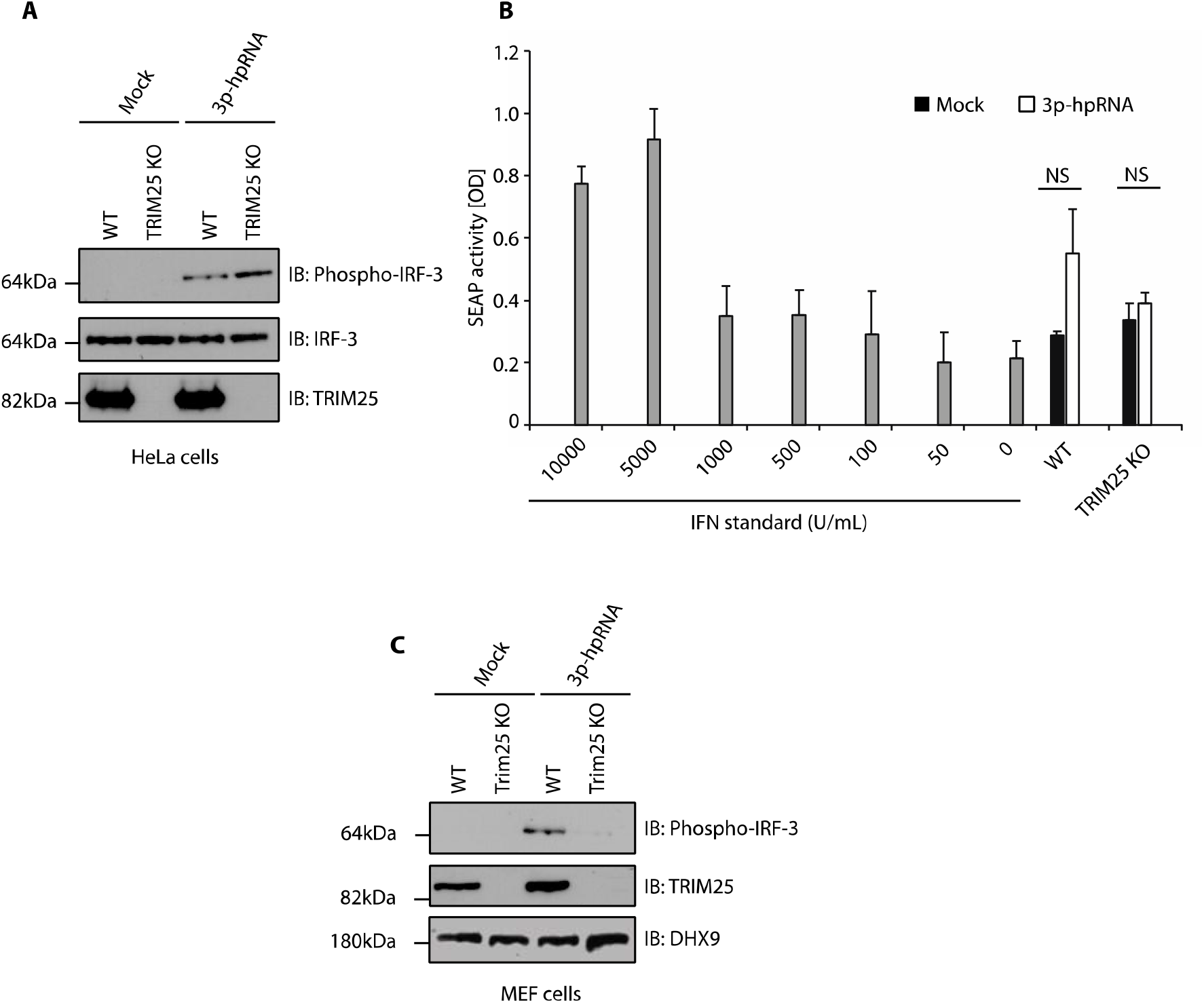
Deletion of TRIM25 in MEF cells, but not HeLa cells, reduces activation of RIG-I. (A) Western blot analysis of IRF-3 phosphorylation after 6 hours in HeLa WT and TRIM25 KO cells transfected with 100 ng/mL 3p-hpRNA. (B) HEK-Blue assay of supernatant from HeLa cells 6 hours post-transfection with 100 ng/mL 3p-hpRNA. The means and SDs of three independent experiments are shown. Statistical significance was calculated using Welch’s t-test. (C) Western blot analysis of IRF-3 phosphorylation after 6 hours in MEF WT and TRIM25 KO cells transfected with 100 ng/mL 3p-hpRNA.

Overall, these results suggest that RIG-I signalling can function efficiently in human HEK293 and HeLa but not mouse MEF cells in the absence of TRIM25. Thus, our results confirmed the dependence on TRIM25 in RIG-I/interferon type I activation in mouse cells ^5^ while provided further evidence that TRIM25 activity towards the RIG-I pathway is redundant in selected human cultured cells. There is likely more variation between different species and even between cell lines of the same species as the role of TRIM25 may be defined by the presence of other proteins and cellular factors.

### The presence of TRIM25 does not restrict transcription of viral mRNAs

A previous study has shown that TRIM25 can inhibit transcription of IAV RNAs by blocking IAV’s polymerase ^29^. To test if this occurred in our cells, we performed a viral minireplicon assay. This involves transfecting cells with constructs expressing the components required for viral mRNA synthesis (polymerase subunits (PA, PB1, PB2) and nucleoprotein NP) as well as a reporter plasmid encoding Firefly luciferase. The host machinery transcribes the Firefly luciferase gene as a negative sense RNA flanked by the 5’ and 3’ sequences of an IAV vRNA. For Firefly luciferase protein to be expressed, this negative sense RNA must be transcribed to a positive sense mRNA, a process that requires an active IAV RNA polymerase and therefore any inhibition of viral mRNA synthesis by TRIM25 can be measured. This assay was performed in the HEK293 cell lines and Firefly luciferase activity was determined (Fig. 5A). Luciferase activity was not significantly different between WT, TRIM25 KO, TRIM25 KO + TRIM25ΔRBD or TRIM25 KO + TRIM25ΔRING cell lines, indicating that IAV RNA polymerase activity was not inhibited by the presence of TRIM25, at least for this luciferase reporter gene. Importantly, dynamine-like large GTPase MX1, a known inhibitor of IAV polymerase ^39^, efficiently inhibited luciferase production, validating the negative results obtained in TRIM25 KO cells.

**Figure 5.**
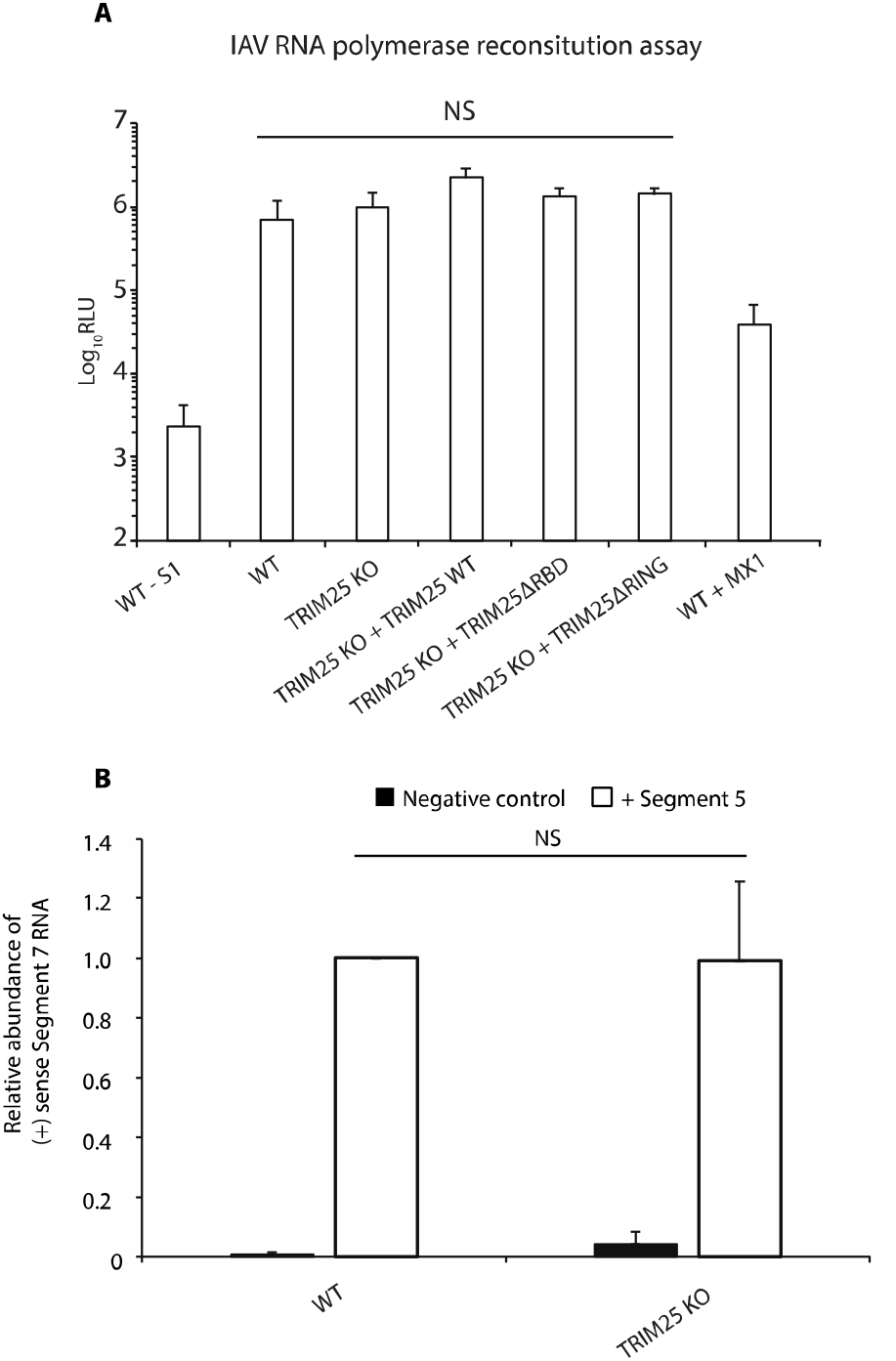
IAV RNA polymerase activity is insensitive to TRIM25 in HEK293 cells. (A) The indicated HEK cell lines were transfected with plasmids to reconstitute IAV RNPs as well as a reported plasmid encoding Firefly luciferase that is transcribed in the negative sense. Negative control experiments were performed in the same way but segment 5 (encoding NP) of the IAV genome was omitted, preventing IAV RNA polymerase activity. The means and SDs of three independent experiments are shown. Statistical significance was calculated using one-way ANOVA with Tukey’s post-test. (B) IAV RNPs were reconstituted along with segment 7 vRNA. Levels of positive sense segment 7 RNA produced by the IAV polymerase were assayed by qRT-PCR and normalised to levels of GAPDH mRNA. The means and SDs of three independent experiments are shown. Statistical significance was calculated using Welch’s t-test.

To further test the effect of modulating TRIM25 expression, the assay was performed with segment 7 of the IAV genome, encoding M1 and M2 proteins, as a reporter to see if TRIM25 inhibited IAV RNA polymerase activity on an authentic genome segment. The assay was performed in the same way as previously but transcription from negative to positive-sense mRNA and cRNA was analysed by qRT-PCR detecting positive sense RNA from segment 7 (Fig. 5B). Negative control experiments performed without NP produced only low levels of RNA, as expected ^40^, whereas reconstitution of complete RNPs produced more than 100-fold more RNA. Once again, however, no difference was seen between WT and ΔTRIM25 cells, indicating that TRIM25 was not inhibiting IAV RNA polymerase activity. Taken together, these results suggested that TRIM25 is not inhibiting IAV gene expression in HEK293 cells, despite substantially inhibiting overall replication of the IAV PR8 (NS1 R38K41A) strain (Fig. 1B). In summary, our results suggest a new yet unidentified mechanism that TRIM25 uses to inhibit IAV infection.

### TRIM25 destabilises IAV mRNAs during infection

Since TRIM25 binds viral positive strand RNAs and restricts IAV PR8 infection, we wanted to investigate if this binding influenced the stability of the IAV RNAs. We treated HEK293 WT and HEK293ΔTRIM25 IAV PR8 infected cells with Favipiravir (T-705), a drug that inhibits the viral polymerase ^41^, and measured the changes in the amount of viral RNA segments, using qRT-PCR, in the cells at different time points. This showed that following Favipiravir treatment the viral RNAs were more stable in HEKΔTRIM25 cells than in HEK293 WT cells (Fig. 6). While in control (DMSO-treated cells) there was no difference, with two-way ANOVA and Sidak’s post-test showing significant difference in T-705 treated cells compared to DMSO (Fig. 6). Crucially, qRT-PCR does not differentiate between functional/non-functional or intact/partially degraded RNAs and thus the lack of difference between WT and TRIM25 KO cells for DMSO-treated cells might not be showing the full picture of TRIM25 action in cells.

**Figure 6.**
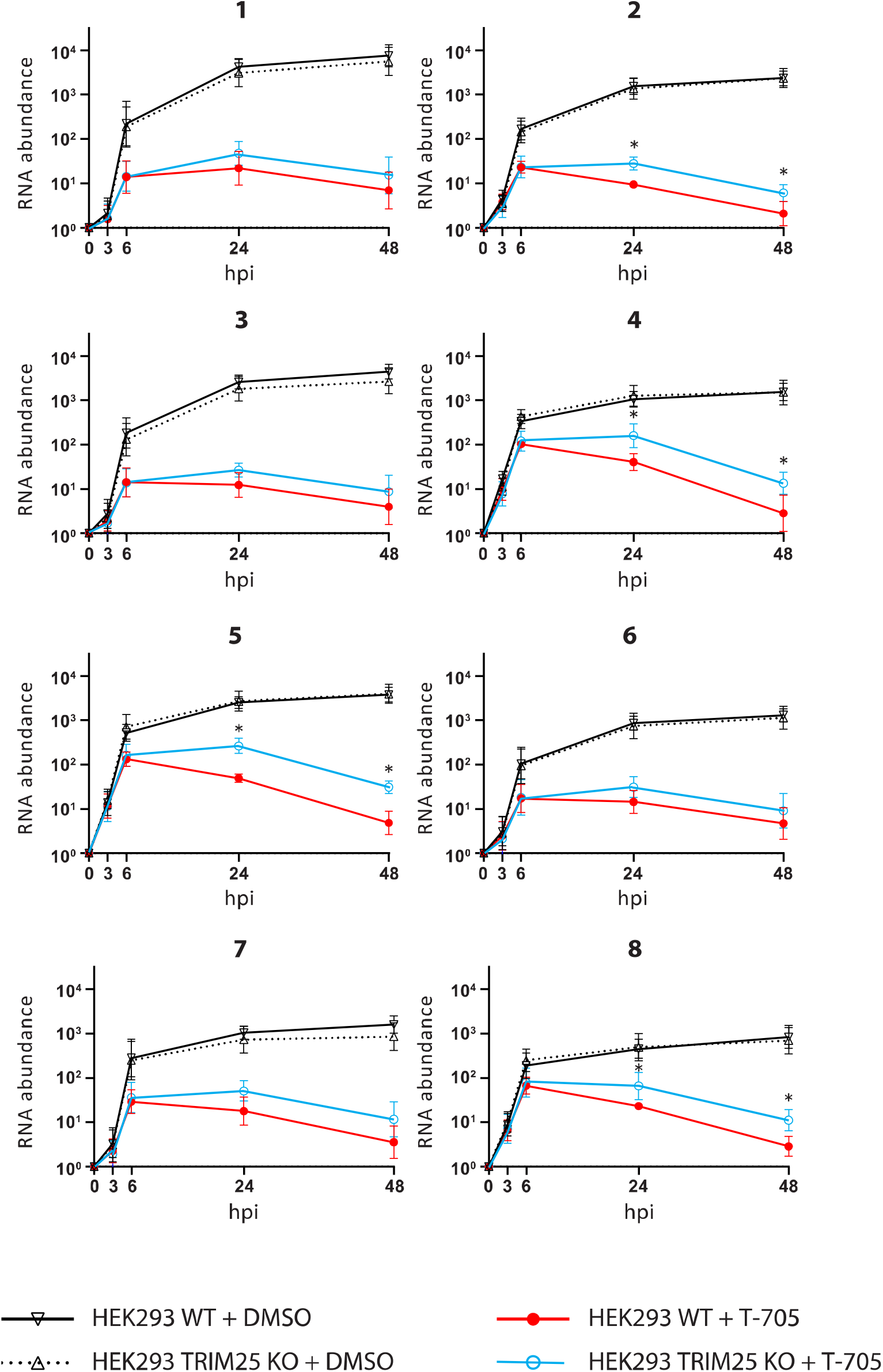
TRIM25 regulates stability of IAV RNA. HEK293 WT and HEK293ΔTRIM25 were infected with IAV PR8 R38K41A and were treated with Favipiravir (T-705) or DMSO (control). The relative changes in the amount of viral RNA segments (indicated with a number over the graphs) were measured using qRT-PCR in the cells at different time points. The level of RNA at 0 hpi was adjusted to one. Statistical significance was calculated using two-way ANOVA with Sidak’s post-test upon log-transformation. Treatment of IAV infected cells with T-705 gave statistically significant results at 24 and 48 hpi in HEK293ΔTRIM25 compared to HEK293 WT cells.

Since a standard qRT-PCR does not necessarily distinguish between the three different types of IAV RNAs (vRNA, mRNA, cRNA) a qRT-PCR using an oligo dT primer was used for the RT to mainly reverse transcribe mRNAs. Primers for specific viral segments were then used for the qPCR. This showed that at 48h post treatment six out of eight segment-derived mRNAs were significantly decreased in the presence of Favipiravir in WT cells (Fig. 7) when compared to HEK293 ΔTRIM25-Favipiravir treated cells. These results show that TRIM25 binds to IAV mRNAs leading to their destabilisation.

**Figure 7.**
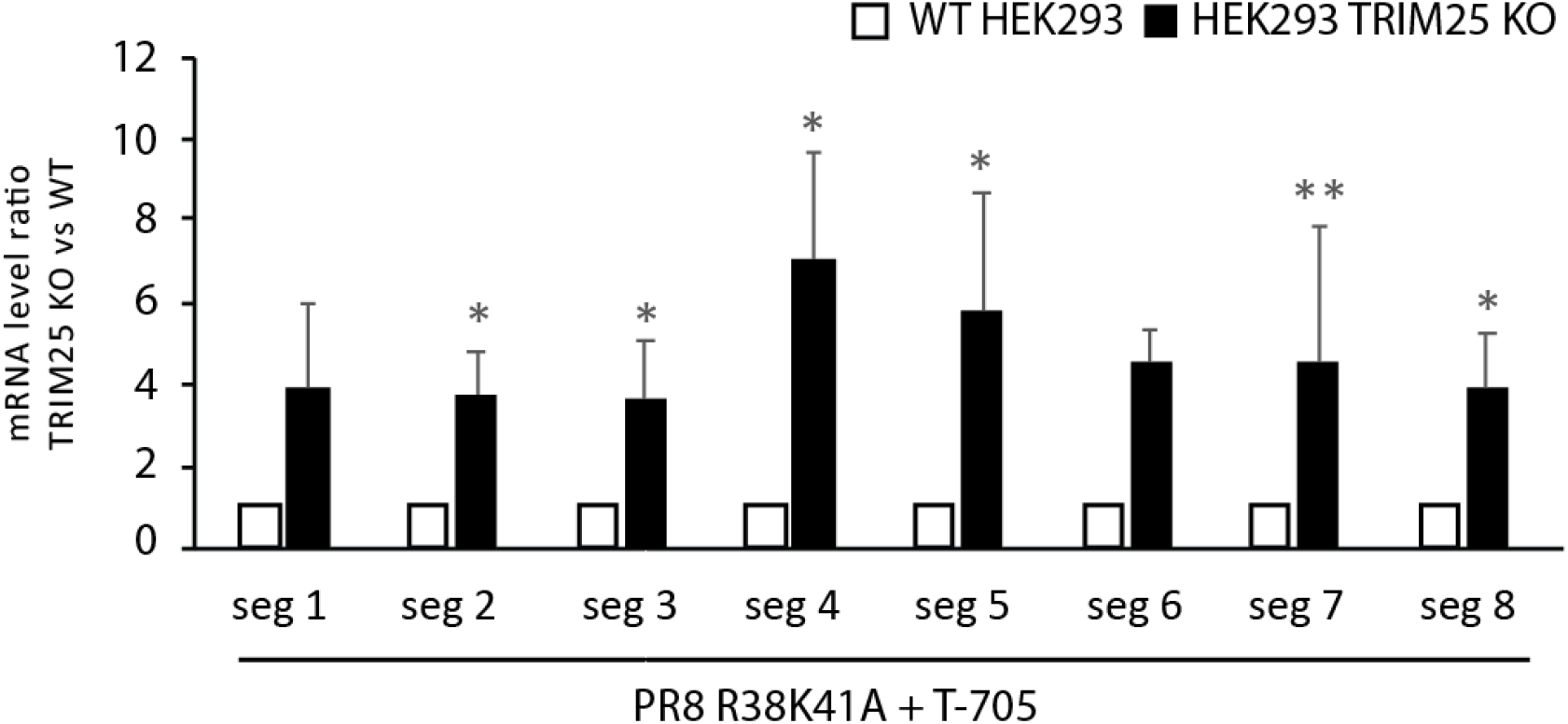
TRIM25 regulates stability of IAV mRNA. HEK293 WT and HEK293ΔTRIM25 were infected with IAV PR8 R38K41A and were treated with Favipiravir (T-705) or DMSO control. The changes in the amount of viral mRNA segments were measured using qRT-PCR in the cells at different time points. To quantify viral mRNAs a qRT-PCR with an oligo dT primer was used for the RT. Primers for specific viral segments were used for the qPCR. The means and SDs of four independent experiments are shown. Statistical significance was calculated using independent Student’s t-test.

### Direct tethering of TRIM25 to mRNA causes RNA degradation

To further investigate TRIM25’s effect on mRNA stability and to test if this is a direct or indirect mechanism, we tethered TRIM25 to Renilla mRNA. To do so, we introduced nine MS2 stem loops into the 3’UTR of Renilla mRNA in a psiCHECK2 plasmid and co-transfected plasmids expressing T7-TRIM25 WT, T7-TRIM25-MS2 fusion protein or MS2 on its own (Fig. 8A-C). Strikingly, only the T7-TRIM25-MS2 fusion protein was able to significantly reduce (P=0.0008) the Renilla luciferase levels as assessed by luciferase assay, and only in the presence of the nine MS2 stem loops (Fig. 8D-E). This provides direct evidence that attracting TRIM25 towards a specific mRNA reduces its expression. To further explore the importance of RNA binding and ubiquitination on TRIM25s ability to destabilise mRNAs we co-transfected T7-TRIM25ΔRBD-MS2 and T7-TRIM25ΔRING-MS2 fusion proteins with the psiCHECK2 constructs. While a slight decrease was seen in Renilla luciferase levels, this was not significant (P=0.62 and 0.99, respectively) as assessed by ANOVA on ranks with Dunnett’s post-test (Fig. 8F-G). This indicated that RNA binding and ubiquitination may be important for TRIM25’s ability to de-stabilise certain mRNAs.

**Figure 8.**
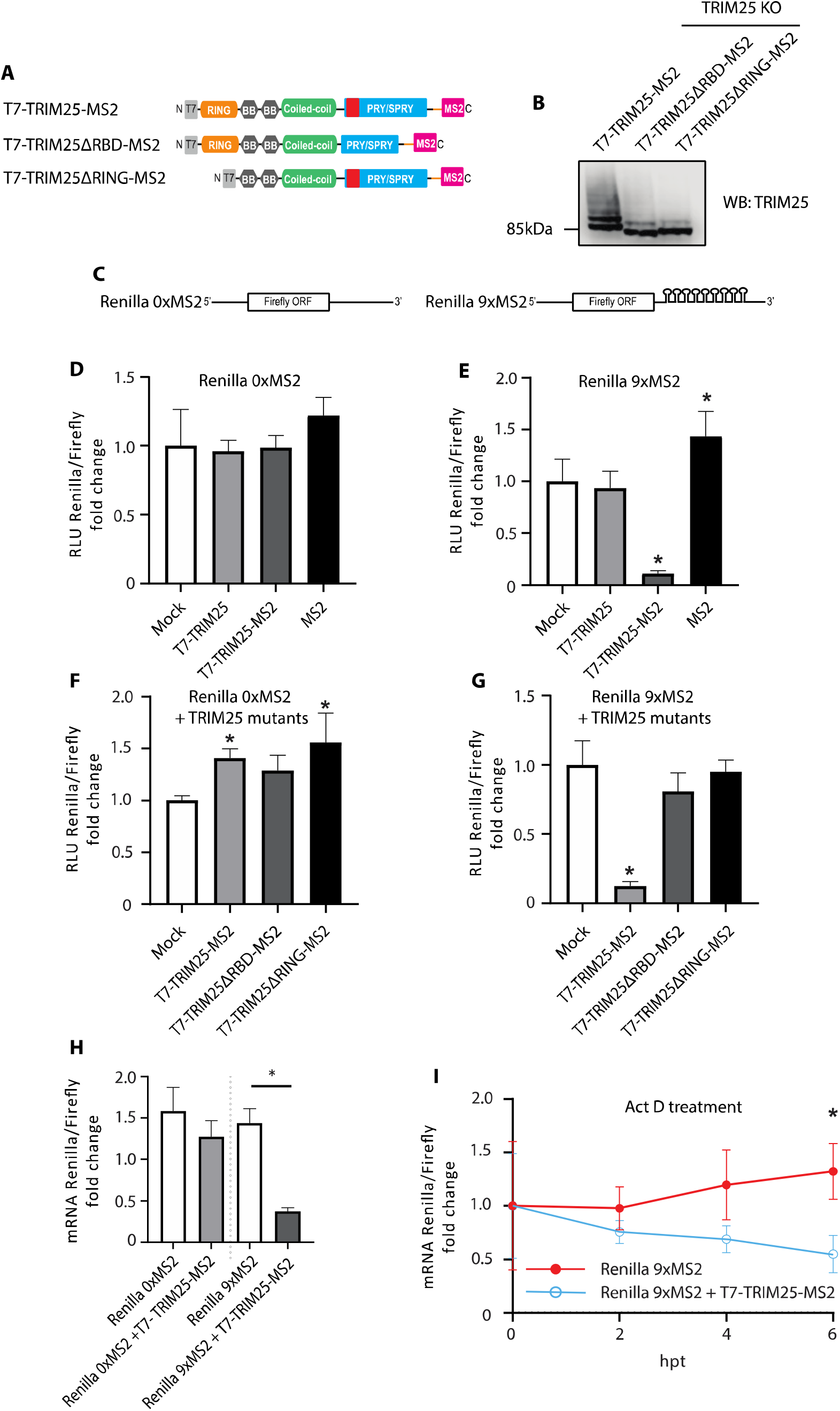
Tethering of TRIM25-MS2 to MS2 stem loops triggers mRNA degradation. (A) Domain architecture of fusion protein of TRIM25, TRIM25ΔRBD or TRIM25ΔRING with MS2 domain (which binds MS2 RNA stem loops), tagged with T7 epitope. (B) T7-TRIM25-MS2, T7-TRIM25ΔRBD-MS2 or T7-TRIM25ΔRING-MS2 were overexpressed in HEK293 TRIM25 KO cells and levels of the proteins were compared to WT cells by western blot. (C) Sequence architecture of Renilla mRNA with and without nine MS2 stem loops located in the 3’UTR. (D-E) The psiCHECK2 plasmids without or with nine MS2 stem loops were co-transfected with plasmids expressing T7-TRIM25 WT, T7-TRIM25-MS2 fusion protein or MS2 on its own in to HEK293 TRIM25 KO cells and analysed with dual luciferase assay. (F-G) T7-TRIM25-MS2, T7-TRIM25ΔRBD-MS2 and T7-TRIM25ΔRING-MS2 fusion proteins were co-transfected with the psiCHECK2 constructs in to HEK293 TRIM25 KO cells and processed with dual luciferase assay. (H) Renilla mRNA levels were assayed upon reverse transcription, followed by DNase treatment and qPCR. Renilla mRNA with and without nine MS2 stem loops was measured after mock or T7-TRIM25-MS2 overexpression. (I) Actinomycin D treatment was preformed after co-expressing the psiCHECK2 plasmid containing nine MS2 stem loops with T7-TRIM25-MS2 fusion protein. Renilla mRNA levels were measured upon reverse transcription, followed by DNase treatment and qPCR at 0, 2h, 4h, and 6h following actinomycin D treatment. The means and SDs of three (D, E, H, I) or six (F, G) independent experiments are shown. Statistical significance was calculated using one-way ANOVA with Dunnett’s (D, E) or Sidak’s (H) post-test, two-way ANOVA with Sidak’s post-test (I) and ANOVA on ranks (F, G) The asterisk (*) indicates p < 0.05.

To confirm that TRIM25 influences Renilla mRNA levels when tethered to it, we measured Renilla mRNA levels by qRT-PCR after DNase treatment of the RNA (Fig. 8H). Importantly, we saw a significant decrease in Renilla mRNA levels only when co-expressing the psiCHECK2 construct containing nine MS2 stem loops and T7-TRIM25-MS2 fusion protein confirming that TRIM25 acts on mRNA level. Actinomycin D treatment confirmed that TRIM25 affects mRNA stability, since only the Renilla mRNA of the construct with nine MS2 stem loops was affected by actinomycin D treatment when co-expressed with T7-TRIM25-MS2 fusion protein (Fig. 8I).

## Conclusions

Here, we show that deletion of TRIM25 from HEK293 cells relieved inhibition of an NS1 mutant of IAV unable to fully control innate immune responses. As would be expected, inhibition was restored by re-integration of TRIM25 WT, but surprisingly, the host RNA binding-deficient mutant TRIM25ΔRBD and the ubiquitination activity-deficient TRIM25ΔRING also rescued the TRIM25 KO phenotype. This suggests that TRIM25 may perform another role in restricting IAV not involving its E3 ubiquitin ligase activity. Interestingly, our CLIP-seq results revealed that that TRIM25ΔRBD could still bind viral RNAs, using a yet unknown mechanism, indicating that RNA binding is important for TRIM25’s function. We also found that deletion of TRIM25 did not significantly affect the RIG-I response to an IAV-derived 5’ppp-RNA in these same cells, contrary to previously described results ^6,42^. This is in line with reports showing that RIPLET could be more important for RIG-I activation in human cells ^14,15^. RIG-I response is attenuated by TRIM25 knockout in MEF cells, showing that the role and necessity of TRIM25 may vary in the context of different cell types and organisms. We further show that human TRIM25 does not seem to directly inhibit transcription of vRNAs in HEK293 cells, as has been shown in a previous study ^29^. Here for the first time, we show that TRIM25 binding to viral mRNAs triggers their degradation (Fig. 9). These observations were reinforced with the RNA tethering experiment where TRIM25 directly inhibited levels of bound mRNA. We note that RNA stability assays with MS2 stem loops, unlike IAV infections indicated the need for RING and RBD domains. We thus speculate that TRIM25 might have other RING and RBD-independent functions that are crucial for its role in inhibiting viral infection.

**Figure 9.**
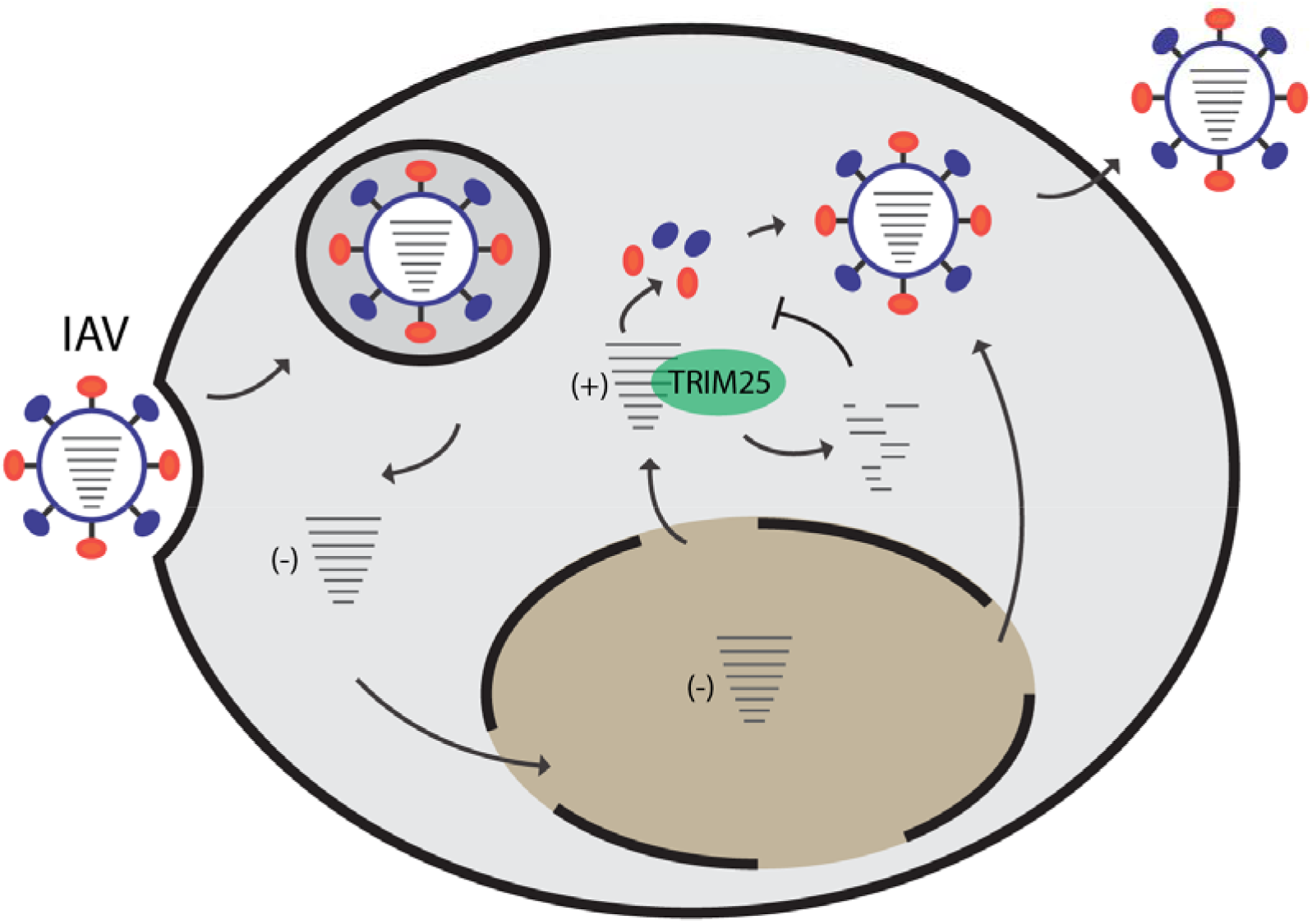
Model of TRIM25 sensing and inhibiting IAV infection by controlling viral mRNA stability. During IAV infection TRIM25 bind to abundantly produced positive strand RNAs and triggers their degradation, which in turn slow down viral replication. This mechanism is dependent on TRIM25’s E3 ligase activity. We also predict that TRIM25 is involved in other, E3 ligase-independent mechanisms of controlling IAV infection.

The exact mechanism of TRIM25-mediated degradation of viral mRNAs and its extent to endogenous RNAs or other viruses should be studied next. Nonetheless, TRIM25 is known to interact with the antiviral protein ZAP, which has been shown to degrade viral RNAs (reviewed in ^43,44^). In addition, TRIM25 has been shown to be required for ZAP’s antiviral function, however, the exact mechanism for this remains unclear ^45,46^. It is plausible that ZAP binds to RNA and uses TRIM25 to trigger RNA degradation. Intriguingly, our previous results showed that in undifferentiated mouse cells TRIM25 helps triggering exoribonuclease DIS3L2-mediated degradation of let-7 microRNA precursor^16^. It remains to be established whether similar factors are responsible for TRIM25-mediated degradation of viral mRNAs. Furthermore, while ZAP was found to bind CpG dinucleotides on viral RNAs^47^, our CLIPseq analysis does not reveal this preference for TRIM25 on IAV mRNA, with GpC dinucleotides preferred. Intriguingly, a clear preference for GpC dinucleotide was observed for TRIM25 but not for TRIM25ΔRBD. This could be explained by alternative RNA-binding domains providing broader sequence specificity for TRIM25. Indeed, such domains have been already identified, including 7K motif, situated between coiled coil and the PRY/SPRY domain ^48^. Finally, TRIM25 has been shown to aid ZAP’s RNA binding ability, perhaps to overcome the CpG dinucleotide suppression in viral RNAs. The interactions of TRIM25 and ZAP during viral infection and the effect on the stability of viral mRNAs and virus replication needs to be further investigated.

Our results demonstrate for the first time that TRIM25 is binding IAV mRNA and triggers RNA degradation. This provides additional mechanism by which TRIM25 can restrict IAV infection. It possible that TRIM25’s RNA binding role is important in the context of other RNA viruses. Thus, uncovering the TRIM25 mechanism and protein binding partners used in its RNA degrading role will be of great importance. In summary, our results reveal novel, unexpected features of TRIM25’s roles in innate immune antiviral pathways and could help designing innovative, host-based RNA virus-targeted therapies.

## Materials and methods

### Virus infections

Influenza virus strain A/Puerto Rico/8/1934 H1N1 and its NS1 mutant (R38K41A) were used for infections and were generated by reverse genetics using plasmids encoding all 8 segments of the IAV genome ^32,49^.

To evaluate the viral replication kinetics, HEK293 cells were inoculated in suspension at an MOI of 0.0001 in 300 μL of DMEM supplemented with 0.14% BSA. Cells were then seeded in 12-well plates to obtain a resulting concentration of 2×10^6^ cells per well. Mock-infected cells were included as controls. Cells were incubated with virus (5% CO_2_, 37°C) for 3h before the addition of 2 mL of virus growth medium (DMEM, 0.14% BSA, 0.5 μg/mL TPCK-treated trypsin). Supernatants were collected with three technical replicates every 24h and frozen (−80°C) until subsequent infectious virus quantification with the endpoint dilution assay ^50^. Fifty percent cell culture infective dose (CCID_50_) endpoint titers were calculated by the Spearman–Kärber formula and expressed as a decimal logarithm. Media from mock-inoculated cells were used as negative controls.

For Favipiravir treatments, HEK293 cells were seeded in 6-well plates such that they reached a confluency of 70% before being infected with virus at an MOI of 0.01. Cells were incubated with virus for 1.5 hours before the addition of virus growth media (DMEM, 0.14% BSA, 1 μg/mL TPCK-treated trypsin) supplemented with 500 μM of Favipiravir (Selleck Chemicals) or DMSO. The cells were incubated for indicated times and then cells were resuspended in TRI Reagent (Sigma-Aldrich).

### CLIP-seq

HEK293ΔTRIM25 cells with integrated T7-tagged TRIM25 WT or TRIM25ΔRBD were infected with the IAV PR8 R38K41A strain at an MOI of 5 for 1 or 6 hours followed by UV crosslinking. CLIP-seq was then performed as published previously^21^.

### 3p-hpRNA assays

HEK cell lines were transfected at 70% confluency with 100 ng/mL 3p-hpRNA (Invivogen) using Lipofectamine 2000. For western blots, cells were incubated for 6 hours before protein extraction. 80 μg protein lysate was loaded per well. Proteins were detected using the following antibodies and dilutions: phospho-IRF-3 (Rabbit mAb, Cell Signalling Technologies, 1:1000), IRF-3 (Rabbit mAb, Cell Signalling Technologies, 1:1000), TRIM25 (Rabbit pAb, Abcam, 1:2000), DHX9 (Rabbit pAb, Protein-Tech, 1:1000). For HEK-Blue assays, supernatants were harvested 6 hours post-transfection. 20 μL of the supernatant or an IFNα/β standard containing purified IFNα and IFNβ were added to 180 μL HEK-Blue cells (Invivogen) (50,000 cells) and incubated for 24 hours. 5 μL of this supernatant was added to 180 μL QUANTI-Blue (Invivogen) and incubated for one hour at 37°C before absorbance was read at 680 nm. For luciferase assays, cells were transfected with 100 ng/mL 3p-hpRNA along with a Firefly luciferase reporter under the IFNβ promoter and Renilla luciferase under a constitutive reporter. The cells were incubated for 24 hours post-transfection before extraction with passive lysis buffer (Promega). Activity levels were measured for Firefly and Renilla luciferase using the Dual-Luciferase Reporter Assay System (Promega) according to the manufacturer’s instructions.

### IAV minireplicon assay

Cells were transfected with plasmids encoding segments 1, 2, 3 and 5 (PA, PB1, PB2, NP) of the IAV genome ^32^. For the luciferase reporter assay, a plasmid encoding Firefly luciferase that is transcribed as a negative sense RNA containing flanking sequences derived from IAV, was co-transfected along with the IAV genome segments. Cells were incubated for 48 hours before the luciferase assay proceeded as previously described. For the segment 7 reporter assay, a plasmid encoding IAV segment 7 that is transcribed as a negative sense RNA was co-transfected in addition to the other IAV genome segments. For qRT-PCR experiments, cells were incubated for 48 hours before RNA was extracted using TRI Reagent (Sigma-Aldrich). qRT-PCRs were performed using the SuperScript III Platinum One-Step qRT-PCR kit (ThermoFisher) according to the manufacturer’s instructions. Levels of positive sense segment 7 RNA were measured and normalised to GAPDH using the primers in Table 1.

**Table 1 –.**
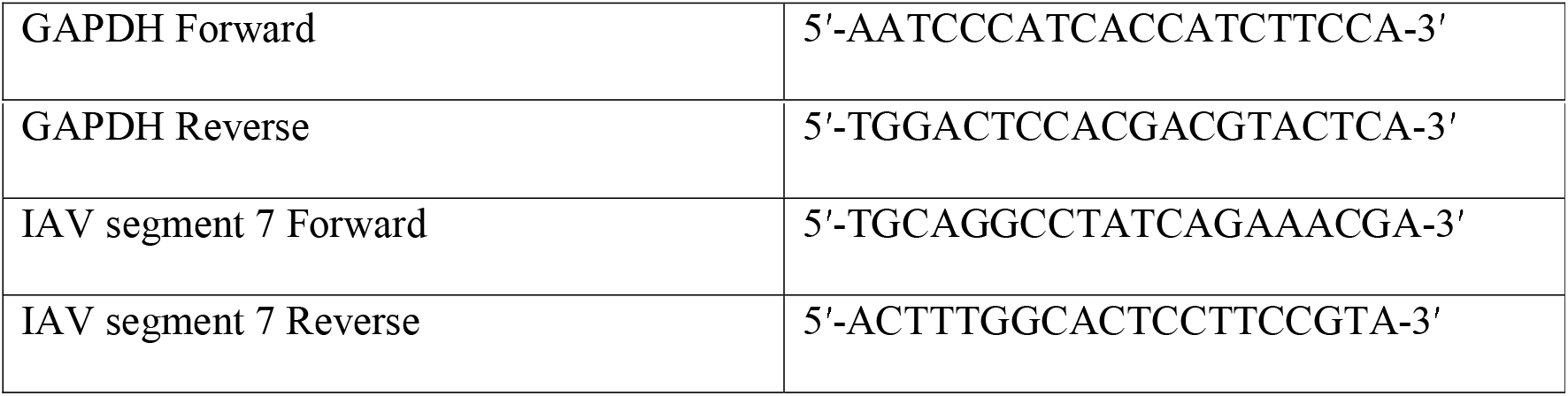
Primers used in qRT-PCR experiments.

#### TRIM25 RNA tethering assays

HEKΔTRIM25 cells were seeded in a 96-well plate at 15,000 cells per well for luciferase assays, or in a 24-well plate at 90,000 cells per well for RNA extractions. The following day the plate was transfected with psiCHECK2 containing nine MS2 stem loops in the 3’UTR of Renilla, or a construct without the nine MS2 stem loops and co-transfected with plasmids as indicated using Lipofectamine 2000 (Invitrogen).

The nine stem loops were cloned in to psiCHECK2 by restriction digest from Luc-MS2 plasmids kindly gifted by Nicola Gray. The psiCHECK2 plasmid provides an internal control of Firefly luciferase, which is used for normalising transfection efficiency and general effects on gene expression.

The TRIM25-MS2 fusion protein was designed by linking the MS2 protein to the C-terminus of TRIM25 with a flexible linker (GSGGGGSRS) between.

After 24 hours a luciferase assay was performed according to the instructions of the Dual-Luciferase Reporter Assay System kit (Promega) or the cells were resuspended in TRI Reagent (Sigma-Aldrich).

For actinomycin D treatment the cells were transfected as above with psiCHECK2 containing nine MS2 stem loops and TRIM25-MS2 fusion protein. The following day actinomycin D was added at a final concentration of 5 μg/ml and cells re-suspended in TRI Reagent at 0h, 2h, 4h and 6h post treatment.

#### qRT-PCR

RNA was extracted according to TRI reagent (Sigma-Aldrich) manufacturer’s instructions. When indicated DNase treatment was performed according to instructions for Turbo DNase (Thermo Fisher Scientific) and a phenol/chloroform extraction performed. Either GoTaq 1-Step RT-qPCR System (Promega) was used or reverse transcription for mRNAs was performed using Superscript IV RT (Thermo Fisher Scientific) and oligo(dT) (Thermo Fisher Scientific) and qPCR using GoTaq qPCR Master Mix (Promega) on a qTower (Analytik Jena).

#### Statistical Analysis

All statistical tests applied are indicated in figures caption, as well as the number of samples analyzed. Data were expressed as mean ± standard deviation (M ± SD). A p-value□<□0.05 was considered statistically significant.

## Authors’ contributions

G.M. conceived this study; G.M., P.D., N.R.C., I.T. and G.H. designed the experiments; N.R.C., G.H., I.T., R.M.D.S.P and E.G. performed experiments; GM, P.D., N.R.C., G.H, I.T., R.M.D.S.P and E.G. analyzed data; M.T. designed and prepared TRIM25-MS2 construct; G.M., N.R.C. and G.H. wrote the manuscript with input from other authors. All authors read and approved the final manuscript.

---

**Supplementary Figure 1.**
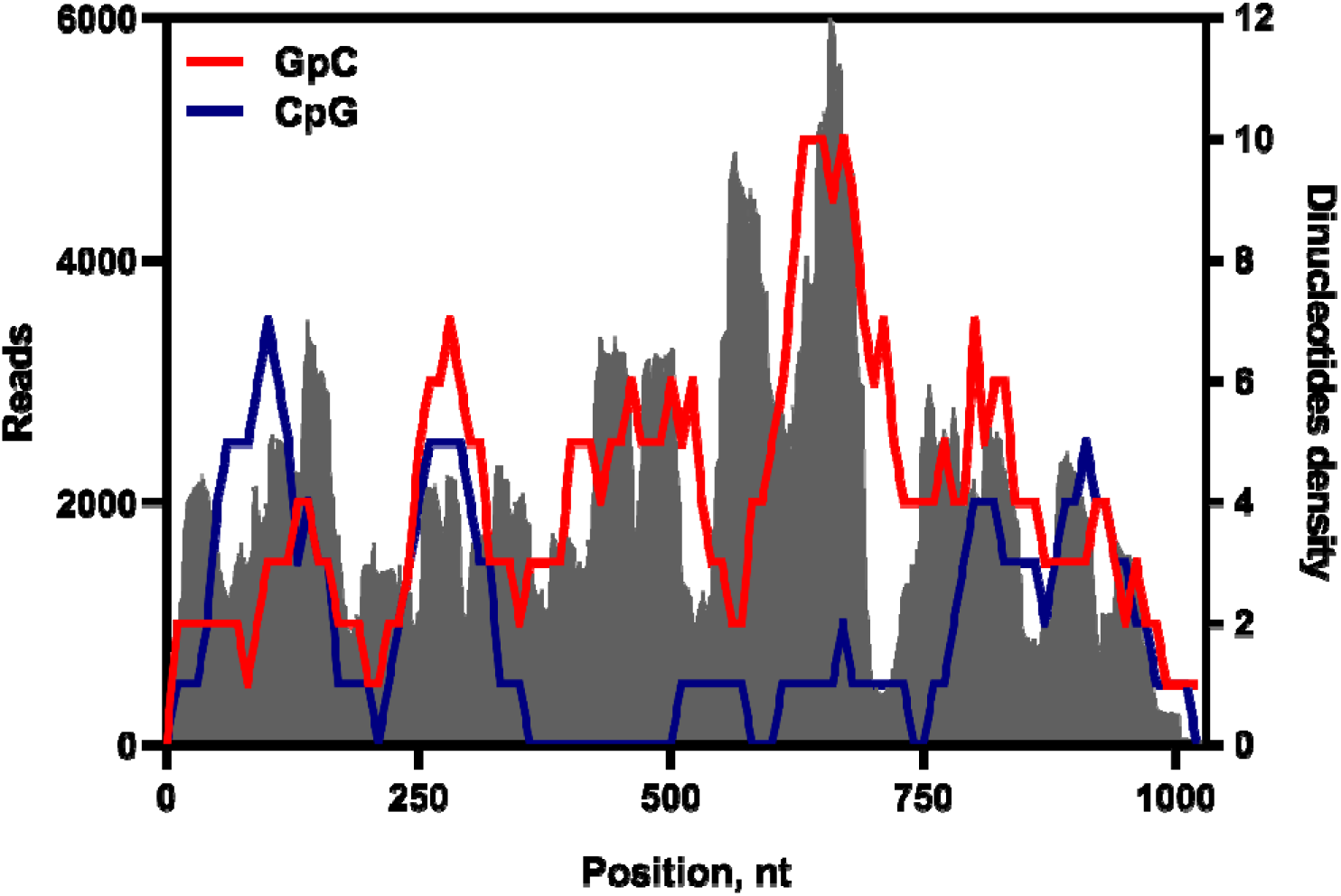
Correlation of mapped reads with specific dinucleotide density. Grey shaded area represents absolute number (left Y-axis) of mapped in CLIP-seq reads for segment 7 of the IAV genome at 6 hpi in HEK WT TRIM25 cells. An R script (https://github.com/itrus/Mapping-ROIs-in-genetic-sequences) was used to count dinucleotides in a 70-nucleotide sliding window. Specific dinucleotide frequencies are represented with lines (right Y-axis).

